# Comprehensive Assessment of the Intrinsic Pancreatic Microbiome

**DOI:** 10.1101/2023.08.12.553074

**Authors:** Austin M. Eckhoff, Ashley A. Fletcher, Matthew S. Kelly, Anders Dohlman, Caitlin A. McIntyre, Xiling Shen, Matthew K. Iyer, Daniel P. Nussbaum, Peter J. Allen

## Abstract

**Background:** Small studies in pancreatic ductal adenocarcinoma (PDAC) and intraductal papillary mucinous neoplasm (IPMN) have suggested that intra-pancreatic microbial dysbiosis may drive malignant transformation. We sought to comprehensively profile tissue and cyst fluid in patients with benign, precancerous, and cancerous conditions of the pancreas to characterize the intrinsic pancreatic microbiome.

**Methods:** Pancreatic samples were collected at the time of resection from 109 patients. Samples included tumor tissue (control, n=20; IPMN, n=20; PDAC, n=19) and pancreatic cyst fluid (IPMN, n=30; SCA, n=10; MCN, n=10). Assessment of bacterial DNA by quantitative PCR and 16S ribosomal RNA gene sequencing was performed. Downstream analyses determined the relative abundances of individual taxa between groups and compared intergroup diversity. Whole-genome sequencing data from 140 patients with PDAC in the National Cancer Institute’s Clinical Proteomic Tumor Analysis Consortium (CPTAC) were analyzed to validate findings.

**Results:** Sequencing of pancreatic tissue yielded few microbial reads regardless of diagnosis, and analysis of pancreatic tissue showed no difference in the abundance and composition of bacterial taxa between normal pancreas, IPMN, or PDAC groups. Low-grade dysplasia (LGD) and high-grade dysplasia (HGD) IPMN were characterized by low bacterial abundances with no difference in tissue composition and a slight increase in *Pseudomonas* and *Sediminibacterium* in HGD cyst fluid. Decontamination analysis using the CPTAC database confirmed a low-biomass, low-diversity intrinsic pancreatic microbiome that did not differ by pathology.

**Conclusions:** Our analysis of the pancreatic microbiome demonstrated very low intrinsic biomass that is relatively conserved across diverse neoplastic conditions and thus unlikely to drive malignant transformation.

**Significance of this study:** *What is already known on this subject?:*

- Microbial colonization, infection, and dysbiosis have been implicated in the oncogenesis of various gastrointestinal tumors (e.g.: *Helicobacter pylori* in gastric cancer, hepatitis B and C in hepatocellular carcinoma, colon dysbiosis in colon cancer progression).
- Few studies have analyzed the intrinsic pancreatic microbiome, and these have produced conflicting results regarding microbial presence and alterations associated with malignant disease.
- IPMN is a precursor lesion to pancreatic cancer that represents a whole gland defect without an established driver event, and microbiome changes have been implicated as a possible etiology of cyst formation and dysplastic progression.

*What are the new findings?:*

- A low-biomass, low-diversity intrinsic pancreatic microbiome is present in both pancreatic tissue and cyst fluid.
- This intrinsic pancreatic microbiome does not differ in terms of abundance, composition, or diversity between patients with PDAC, IPMN, or other benign conditions of the pancreas.

*How might it impact clinical practice in the foreseeable future?:*

- Microbiome dysbiosis does not appear to be a driver of malignant degeneration of IPMN, and further research is needed to identify drivers of oncogenesis in order for possible chemoprevention strategies to be developed.

## Introduction

Microbial colonization has been implicated in the oncogenesis of various gastrointestinal tumors. [1, 2] Due to the pancreas’ direct connection to the gastrointestinal tract, it has been proposed that microbial dysbiosis may drive the development of pancreatic neoplasia, including pancreatic ductal adenocarcinoma (PDAC). [3] However, few prior studies have analyzed the intrinsic pancreatic microbiome, and these have shown discordant results, making it unclear if an intrinsic pancreatic microbiome exists and if these microbial communities drive pancreatic oncogenesis. [3–8] Moreover, the microbiome field is constantly evolving with new technologies and computational methods to optimize detection and contamination. In fact, our group recently analyzed the raw sequencing data from a high-profile publication linking PDAC and microbiome colonization and were unable to reproduce the results, further sparking our interest in this field.[9]

Previous microbiome work has mainly focused on PDAC. However, intraductal papillary mucinous neoplasm (IPMN) is a cystic precursor lesion of pancreatic cancer that represents a “field defect” in that the entire gland is at risk for malignancy, yet no specific germline alteration or other drivers of oncogenesis have been established. [10] We hypothesized that this field defect could be secondary to an abnormal microbial presence, such as seen in gastric cancer and hepatocellular cancer, where pathogenic bacterial (i.e., *H. pylori*) and viral (i.e., hepatitis B/C) infection, respectively, have been implicated in whole-organ risk for malignant progression. [11, 12] This prompted our group to ask whether IPMN dysplasia may also be driven by microbiome alterations which then correlate with malignant transformation from low-grade (LGD) to high-grade (HGD) dysplasia and ultimately to pancreatic cancer.

We sought to comprehensively characterize the intrinsic microbiome of pancreatic tissue and cyst fluid from patients with IPMN and PDAC, as well as the following control samples: normal pancreatic tissue from patients with pancreatic neuroendocrine tumors (PNET), mucinous cystic neoplasms (MCN), and serous cystadenomas (SCA). Our goal was to evaluate evidence of an intrinsic pancreatic microbiome and to determine if these microbial communities are associated with malignant degeneration.

## Methods

### Study population and sample collection

We analyzed samples from 109 subjects enrolled in a biobanking protocol at Memorial Sloan Kettering (MSK) and Duke University Health System (DUHS) between January 2019 and March 2020. Subjects were eligible if they had a PDAC, PNET, IPMN, MCN, or SCA diagnosis. Written informed consent was obtained prior to surgery, and portions of resected tissue and cyst fluid were banked at the time of resection for research purposes.

At DUHS, a protocol to reduce the risk of contamination was established for tissue processing. Samples were collected with sterile instrumentation, snap-frozen in liquid nitrogen, and transferred to a -80°C freezer within one hour of resection. Sample collection at MSK had a similar collection protocol with two exceptions. Some cyst fluid samples from MSK were obtained through endoscopic fine needle aspiration (FNA) and all MSK tissue samples were frozen in optimal cutting temperature compound (OCT). A pancreatic pathologist confirmed tissue diagnosis by separate analysis of tissue sections.

For disease-free control tissue, we obtained histologically normal pancreatic tissue acquired during resection from patients with small, low-grade PNETs. To control for contamination that might be introduced during tissue procurement or DNA extraction, four cryovials of sterile PBS were opened in both the OR and surgical pathology suite and processed in the same fashion as our test samples. Three saliva samples were used as positive controls for DNA extraction.

### DNA Extraction

Total genomic DNA was extracted from human tissue and cyst fluid using the QIAamp Mini kit (Qiagen, Hilden, Germany). Extractions were performed per manufacturer’s instructions, except that samples received overnight treatment with proteinase K at 55 °C, and tissue was mechanically disrupted by beads to further lyse bacterial cell walls (VWR Mini Bead Mill).

### 16S Ribosomal RNA Gene Sequencing

16S-EZ ribosomal RNA (rRNA) library preparations and sequencing on an Illumina MiSeq instrument (Illumina, San Diego, CA, USA) were conducted at GENEWIZ, Inc. (South Plainfield, NJ, USA). The sequencing library was prepared using a MetaVx™ Library Preparation kit (GENEWIZ, Inc., South Plainfield, NJ, USA). V3 and V4 regions were amplified using forward primers containing the sequence ’CCTACGGRRBGCASCAGKVRVGAAT’ and reverse primers containing the sequence ’GGACTACNVGGGTWTCTAATCC’. Five additional PCR cycles were added to the standard protocol to increase the sensitivity of detecting bacterial reads in a potentially low biomass environment. Indexed adapters were added to the ends of the 16S rRNA gene amplicons by limited cycle PCR. DNA libraries were validated and quantified before loading. The pooled DNA libraries were sequenced using a 2x250 paired-end (PE) configuration. Raw sequence data (.bcl files) were converted into FASTQ files and de-multiplexed using Illumina’s bcl2fastq 2.17 software.

### 16S qPCR

Leftover DNA from the sequencing run was sent to the University of North Carolina’s Microbiome Core Facility for 16S quantitative PCR (qPCR) using the same V3/V4 primers used for sequencing. 2 μl of each DNA sample (0.5 ng total) was added to each reaction mix with 2.5 nM final primer concentration for both the forward and reverse primers. Thermo Fisher PowerSybr qPCR master mix was used in each reaction, and samples were analyzed on the QuantStudio6 RealTime PCR instrument. Enzyme activation was performed by incubating each sample at 55°C for 5 minutes, followed by 95°C for 10 minutes. Post activation, samples were cycled for 40 cycles of denaturing at 95°C followed by annealing and elongation at 60°C. Following each elongation step, each well was imaged for Sybr activation. Following the final measurement, melt curve analysis was run on each sample to assess amplicon purity and to identify off-target amplification. 16S rDNA amplicon standard curve from 108 copies to 102 copies was used to calculate 16S rDNA gene copy number in each sample.

### Taxonomic Assignment Pipeline of 16S rRNA Amplicon Sequencing Data

The resulting paired-end FASTQ files were run through the Dada2 pipeline using default parameters, with two maximum errors allowed (maxEE) for both forward and reverse reads. [13] Chimeras were removed using Dada2’s removeBimeraDenovo function, and taxonomy was assigned using DECIPHER (v2.20.0) and the Genome Taxonomy Database (GTDB) training set (vR202). The annotated taxa were run through the decontam (v1.18.0) pipeline to remove possible contaminants based on the initial DNA quantification (frequency) and prevalence in negative control samples using a threshold of 0.5. [14] Sequences that did not align to the GTBD database at the genus level were blasted against NCBI’s nucleotide database and reassigned to the corresponding ASV if the percent identity was ≥ 97% in one of the top 5 hits. Taxa that were not resolved beyond the domain level were removed from downstream analysis. [15]

### Statistical Analysis of qPCR data

Bacterial abundances were estimated by multiplying the copy number/µL of DNA returned by qPCR analysis by the number of µL loaded into each sample’s library prep. Differences in abundance between samples were calculated in RStudio using the Kruskal-Wallis test with FDR correction followed by Dunn’s test. P-values ≤ 0.05 were statistically significant.

### Statistical Analysis of 16S NGS data

For downstream analysis, taxa were filtered from 109 samples using a prevalence threshold of 0.02 and an abundance threshold of 5. Differentially abundant taxa were determined using Wilcoxon’s rank sum test using FDR correction on the top relative abundant taxa at the phylum (top 5) and genus (top 10) levels. Alpha and phylogenetic diversity were compared using Wilcoxon’s rank sum test, and beta diversity was compared using PERMANOVA. All statistical tests were two-way analyses.

### CPTAC Data Acquisition and Analysis

All CPTAC sequencing data were accessed from the Genomic Data Commons (GDC) portal in accordance with the CPTAC Data Use Certification Agreement and under the authorization of Duke’s campus institutional review board. We procured data containing whole genome sequencing (WGS), whole exome sequencing (WXS), and RNA sequencing from matched blood, PDAC tumor, and adjacent normal pancreas tissue from 140 patients, along with brain tissue from 99 patients. To identify the microbial composition of these tissues, we used PathSeq to remove human DNA sequences and identify reads of microbial origin. [16] Unambiguously aligned sequencing reads for bacteria at each taxonomic level were aggregated for available WGS data. [16, 17] These were then normalized to reads-per-million (RPM) counts based on the total number of input sequencing reads in each run. Aggregated PathSeq results and associated metadata for each sequencing run were then converted to phyloseq objects for downstream analyses in R. [15]

### Decontamination analysis

The resulting tumor-associated microbial profiles were then screened using a thorough, multi-step decontamination analysis to identify and remove possible contaminants. Bacteria were defined as tissue-resident if they: 1) were detectable in all three sequencing assays (WGS, WXS, and RNA-seq), 2) were found in higher rates in pancreatic tissue than in matched blood samples, 3) were found in higher rates in pancreatic tissue than in sterile brain tissue, 4) were found in both CPTAC and in the 16S rRNA sequencing data from the Duke cohort, and 5) were not associated with any CPTAC sequencing batch.

First, genera not present in all three CPTAC sequencing assays (WGS, WXS, and RNA-seq) were removed. For the remaining 622 genera detected, we used CPTAC WGS data to compare the prevalence of genera in CPTAC tissues from pancreatic cancer patients with (1) their prevalence in matched blood samples and (2) their prevalence in brain tumors derived from another CPTAC cohort. As described in Dohlman et al., genera were classified as potentially tissue-resident if they were more prevalent in pancreatic tissue (p < 0.05) and were detected in fewer than 20% of blood or brain samples, which should theoretically be sterile. [17] Of the resulting 23 genera, we selected 19 that could also be detected in pancreatic tissues from a second PDAC cohort (Duke). Finally, the remaining 19 genera were evaluated to determine if their presence in CPTAC sequencing data could be explained by batch effect, using a Chi-square test to determine if the observed frequency at which genera were detected in each batch was different from expectation (i.e., even representation in each sequencing batch). This removed 7 additional genera, leaving 12 genera whose presence in pancreatic tissues could not be explained by contamination, false-positive alignment, or batch effect.

## Results

### Patient Characteristics

Pancreatic tissue was collected from patients undergoing resection, including 19 patients with PDAC and 20 with IPMN. Normal pancreatic tissue was obtained from 20 patients undergoing resection of PNETs. For cyst fluid analysis, collection was performed either intraoperatively or via FNA on 30 patients with IPMN, 10 patients with MCN, and 10 patients with SCA. Patients with IPMN were evenly split between those with HGD and LGD. We did not note any differences in bacterial abundance and composition in patients who had undergone invasive endoscopy except for a slight increase in *Pseudomonas* in patients with SCA undergoing endoscopy versus those who did not have a prior biopsy or stent (Supplemental Figure 1). Prior antibiotics use within the three months prior to surgical resection did not alter cyst bacteria composition (Supplemental Figure 2). Clinicopathologic factors for all patients are detailed in Table 1.

**Table 1:** Patient Demographics and Clinical Characteristics.

|  | Control Tissue (PNET) | PDAC Tissue | IPMN Tissue | IPMN Cyst Fluid | MCN Cyst Fluid | SCA Cyst Fluid |
| --- | --- | --- | --- | --- | --- | --- |
| N | 20 | 19 | 20 | 30 | 10 | 10 |
| Median Age, years (IQR) | 64 (59-67) | 70 (60-73) | 65 (62-72) | 71 (63-74) | 53 (49-66) | 48 (42-55) |
| Female Sex | 7 (35%) | 9 (47%) | 8 (44%) | 15 (50%) | 10 (100%) | 8 (100%) |
| Male Sex | 8 (40%) | 8 (42%) | 10 (50%) | 15 (50%) | 0 (100%) | 0 (100%) |
| Race |  |  |  |  |  |  |
| Caucasian | 11 (55%) | 12 (63%) | 16 (80%) | 26 (87%) | 8 (80%) | 5 (50%) |
| Black | 2 (10%) | 2 (10%) | 0 (0%) | 0 (0%) | 1 (10%) | 2 (20%) |
| Asian | 1 (5%) | 0 (0%) | 1 (5%) | 1 (3%) | 0 (0%) | 1 (10%) |
| Other/Unidentified | 6 (30%) | 5 (26%) | 3 (15%) | 3 (10%) | 1 (10%) | 2 (20%) |
| Operation |  |  |  |  |  |  |
| Whipple | 11 (55%) | 14 (74%) | 14 (70%) | 11 (37%) | 9 (90%) | 2 (20%) |
| Distal Pancreatectomy | 3 (15%) | 3 (16%) | 3 (15%) | 15 (50%) | 0 (0%) | 6 (60%) |
| EUS/Stent | 15 (75%) | 14 (74%) | 10 (50%) | 16 (53%) | 5 (50%) | 5 (50%) |
| Other | 1 (5%) | 0 (0%) | 1 (5%) | 4 (13%) | 1 (10%) | 0 (0%) |
| Branch Type |  |  |  |  |  |  |
| Main | NA | NA | 7 (35%) | 10 (33%) | NA | NA |
| Side Branch | NA | NA | 6 (30%) | 11 (37%) | NA | NA |
| Mixed | NA | NA | 4 (20%) | 8 (27%) | NA | NA |
| Antibiotics Given at Time of Incision | 15 (100%) | 17 (100%) | 18 (100%) | 30 (100%) | 10 (100%) | 8 (100%) |
| Antibiotics within the Last 3 Months | 5 (33%) | 3 (18%) | 3 (17%) | 5 (17%) | 2 (20%) | 1 (13%) |

### Pancreatic Tissue Analysis by Disease State (Normal, IPMN, PDAC)

Analysis of 16S qPCR identified a low-density bacterial community within all tissue samples, and overall abundance did not differ by disease state (Figure 1). There was no statistically significant difference in the estimated bacterial DNA abundance between normal pancreas, IPMN tissue, and PDAC tissue (5,518 16S copies/ng DNA Normal vs 3,706 16S copies/ng DNA IPMN vs 3,788 16S copies/ng DNA PDAC; p = 0.06).

**Figure 1:**
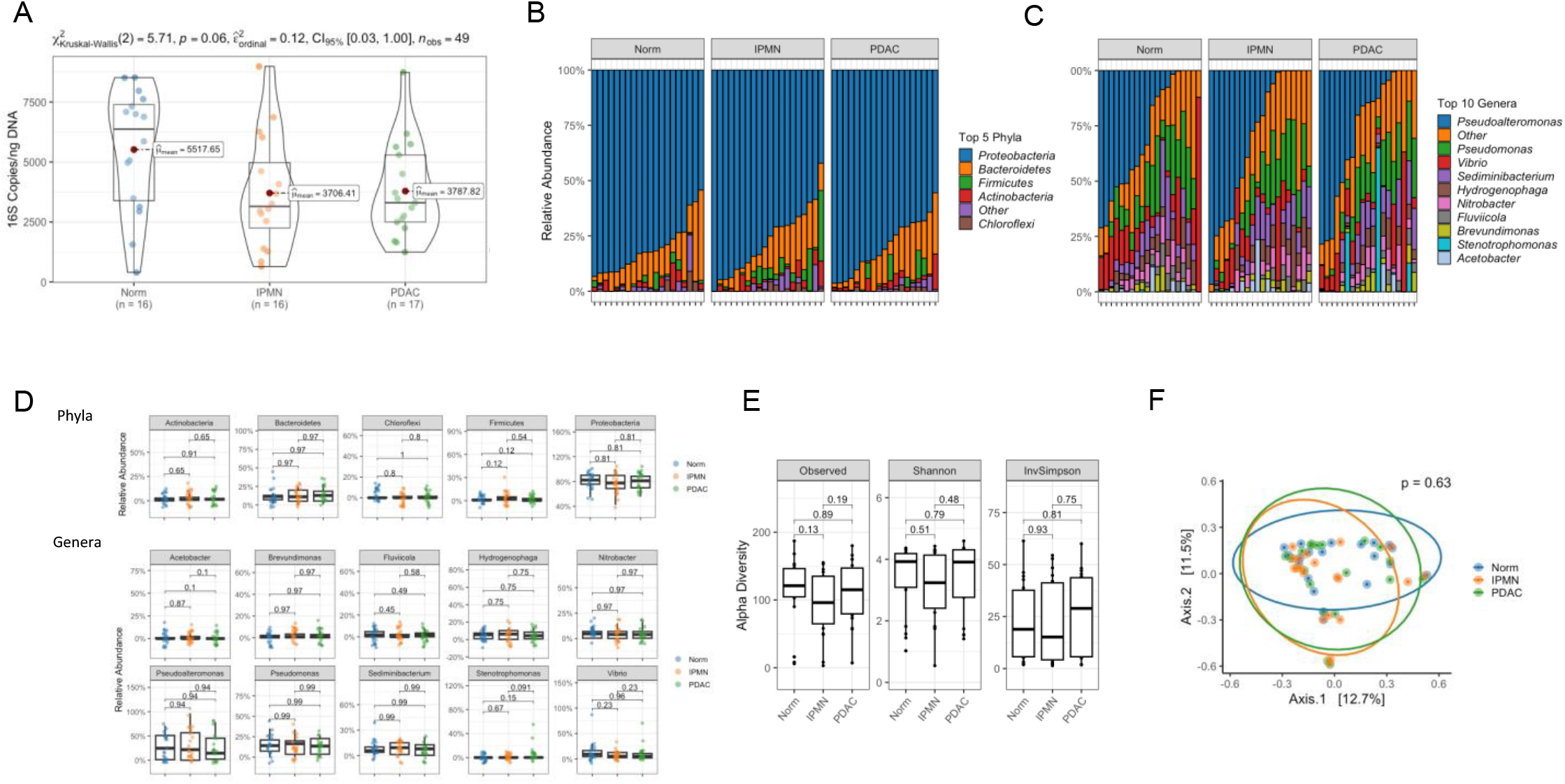
Comparing Pancreatic Microbiome Across Tissue Types (Normal, IPMN, PDAC) A) Absolute read counts of 16S copies/ng of DNA as determined by qPCR. The mean is marked with a red dot, and the median with a horizontal bar. No statistically significant difference was found in the 16S copies/ng DNA between normal pancreas, IPMN, and PDAC tissue as determined by a Krustal-Wallis test with FDR correction. B) Relative abundances of the most abundant bacterial phyla identified in normal pancreas, IPMN, and PDAC tissue. C) Composition plots of the most abundant bacterial genera, based on relative abundance, identified in normal pancreas, IPMN, and PDAC tissue. D) Box plots of relative abundance showing the top phyla and genera between normal pancreas, IPMN, and PDAC tissue. E) Alpha diversity as measured by Observed ASVs, Shannon, and InvSimpson indices between normal pancreas, IPMN, and PDAC tissue. F) PCoA plot of bacteria communities in normal pancreas, IPMN, and PDAC tissue based on the raw Bray-Curtis dissimilarity matrix. Box plot minima and maxima bounds represent the 25^th^ and 75^th^ percentiles, respectively; the center bound represents the median (**D, E**). Whiskers extend to 1.5 times the interquartile range (IQR) (**D**), and data in (**E**) are presented as mean ± SEM. P values were estimated using two-sided Wilcoxon rank-sum tests with FDR correction (**D, E**) or two-way PERMANOVA (**F**).

16S rDNA sequencing was used to determine microbial composition, and there was no difference in relative abundance at the phyla or genera level across disease types. The most abundant phyla seen in pancreatic tissue were Proteobacteria (82.3% normal, 77.6% IPMN, 81.1% PDAC), Bacteriodetes (11.4% normal, 11.1% IPMN, 12.8% PDAC), Actinobacteria (0.8% normal, 1.2% IPMN, 1.5% PDAC), and Firmicutes (0.7% normal, 2.3% IPMN, 0.3% PDAC). Similarly, bacterial genera did not differ by disease state, and the top genera (relative abundance) included *Pseudoalteromonas (24.7% normal, 21.9% IPMN, 14.6% PDAC), Pseudomonas (14.0% normal, 16.5% IPMN, 13.5% PDAC), Vibrio (8.9% normal, 5.1% IPMN, 4.9% PDAC),* and *Sediminibacterium (5.6% normal, 9.3% IPMN, 7.9% PDAC).* Additionally, there was no difference in alpha diversity (Observed ASVs, Shannon, and Inverse Simpson indices), or in beta diversity (PCoA based on a Bray-Curtis dissimilarity matrix) (Figure 1).

### IPMN by Grade of Dysplasia

When comparing tissue samples from IPMNs harboring LGD versus HGD, no difference in the estimated bacterial abundance determined by qPCR, composition determined by 16S rDNA sequencing, or alpha and beta diversity was identified (Figure 2). Similarly, there were no differences in estimated abundance in the cyst fluid from IPMNs harboring LGD versus HGD determined by qPCR. We did note small differences in bacteria composition of *Pseudomonas* (0.15% LGD vs 5.1% HGD, p =0.01) and *Sediminibacterium* (0.0% LGD vs 1.3% HGD, p=0.03), which had greater abundance in HGD IPMN cyst fluid. However, these bacteria were present at extremely low levels; for example, Acidobacteria only had a median (interquartile range) of 36 reads/sample. Finally, there was no difference in alpha or beta diversity (Figure 2).

**Figure 2:**
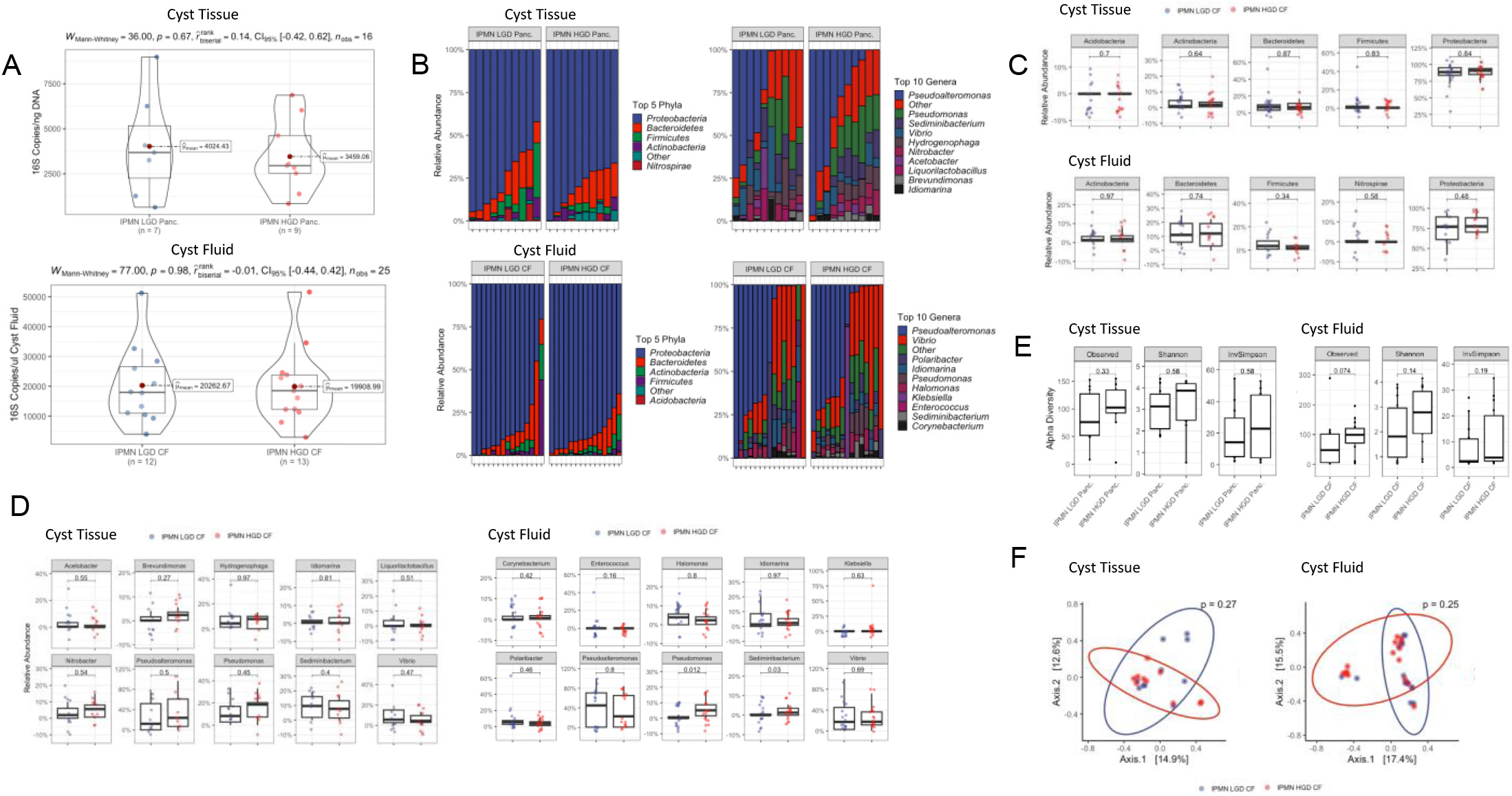
Comparing the Microbiome of Low-Grade Dysplasia vs High Grade Dysplasia IPMN. **A)** Absolute read counts of 16S copies/ng of DNA (left) and 16S copies/ul of cyst fluid (right) as determined by qPCR. The mean is marked with a red dot, and the median with a horizontal bar. No statistically significant difference was found in the 16S copy numbers between the LGD vs HGD IPMN Tissue and LGD vs HGD IPMN cyst fluid as determined by a Mann-Whitney test with FDR adjustment. B) Relative abundances of the most abundant bacterial phyla and genera identified in LGD vs HGD IPMN Tissue and cyst fluid. C) Box plots of relative abundance at the phyla level between the LGD vs HGD IPMN Tissue and LGD vs HGD IPMN cyst fluid. D) Box plots of relative abundance at the genera level between the LGD vs HGD IPMN Tissue and LGD vs HGD IPMN cyst fluid. E) Alpha diversity as measured by Observed ASVs, Shannon, and InvSimpson indices between the LGD vs HGD IPMN Tissue and LGD vs HGD IPMN cyst fluid. F) PCoA plots of bacteria communities in the LGD vs HGD IPMN Tissue (left) and in the LGD vs HGD IPMN cyst fluid (right) based on raw Bray-Curtis dissimilarity matrices. Box plot minima and maxima bounds represent the 25^th^ and 75^th^ percentiles, respectively; the center bound represents the median (**C, D, E**). Whiskers extend to 1.5 times the interquartile range (IQR) (**C, D**), and data in (**E**) are presented as mean ± SEM. P values were estimated using two-sided Wilcoxon rank-sum tests with FDR adjustment (**C, D, E**) or two-way PERMANOVA (**F**).

### Pancreatic Cyst Fluid Analysis by Disease State (IPMN, MCN, SCA)

Analysis of cyst fluid from IPMNs, SCAs, and MCNs demonstrated no differences in bacterial abundance between lesion types based on 16S qPCR. In general, cyst fluid from all three lesions had low bacteria abundance as measured by 16S copies/µl of cyst fluid (20,078 IPMN, 17,305 SCA, 10,938 MCN; p = 0.19).

The five most common bacterial phyla identified within the cyst fluid (relative abundance) from pancreatic cysts were Proteobacteria (90.3% IPMN, 91.8% SCA, 84.0% MCN), Bacteriodetes (6.5% IPMN, 5.1% SCA, 9.8% MCN), Actinobacteria (1.4% IPMN, 1.7% SCA, 2.2% MCN), Firmicutes (0.6% IPMN, 0.9% SCA, 0.6% MCN), and Fusobacteria (0.0% IPMN, 0.1% SCA, 0.3% MCN). Fusobacteria was the only bacterial phylum that differed across cyst types. It was more abundant in MCN than IPMN and present in both at very low levels (p = 0.036). The most common bacterial genera across cyst types (relative abundance) were *Halomonas* (2.5% IPMN, 2.8% SCA, 4.4% MCN), *Idiomarina* (2.3% IPMN, 3.0% SCA, 9.4% MCN), *Polaribacter* (5.0% IPMN, 3.6% SCA, 7.6% MCN), *Pseudoalteromonas* (33.9% IPMN, 63.6% SCA, 0.61% MCN), *Pseudomonas* (1.3% IPMN, 3.3% SCA, 1.8% MCN), and *Vibrio* (17.5% IPMN, 18.8% SCA, 43.8% MCN). *Pseudoalteromonas* was more abundant in SCA than MCN and IPMN (p=0.038).

Finally, there was no difference in alpha or beta diversity between IPMN, SCA, and MCN cyst fluid (Figure 3). Taken together, our tissue and cyst fluid analyses demonstrated that the pancreas has a low biomass intrinsic microbiome for which very limited differences were seen across disease states.

**Figure 3:**
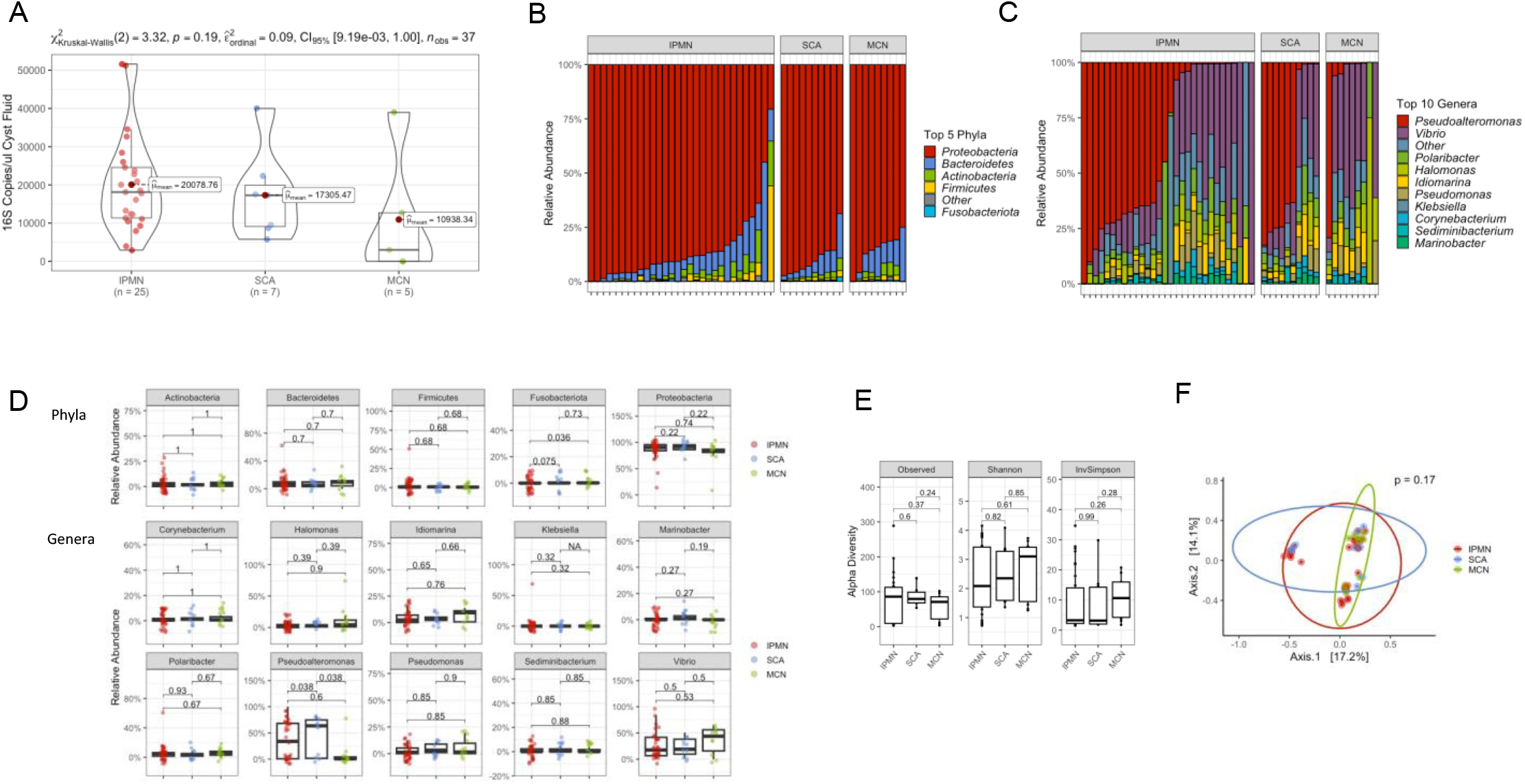
Comparing Cyst Fluid Microbiome Across Diseased States (IPMN, SCA, MCN) A) Absolute read counts of 16S copies/ul of cyst fluid as determined by qPCR. The mean is marked with a red dot, and the median with a horizontal bar. No statistically significant difference was found in the 16S copies/ul of cyst fluid between IPMN, SCA, and MCN cysts as determined by a Krustal-Wallis test. B) Relative abundances of the most abundant bacterial phyla identified in IPMN, SCA, and MCN cysts. C) Relative abundances of the most abundant bacterial genera identified in IPMN, SCA, and MCN cyst fluid. D) Box plots of relative abundance at the phyla and genera level between IPMN, SCA, and MCN cyst fluid. E) Alpha diversity as measured by Observed ASVs, Shannon, and InvSimpson indices between IPMN, SCA, and MCN cyst fluid. F) PCoA plots of bacteria communities in IPMN, SCA, and MCN cyst fluid based on the raw Bray-Curtis dissimilarity matrix. Box plot minima and maxima bounds represent the 25^th^ and 75^th^ percentiles, respectively; the center bound represents the median (**D**). Whiskers extend to 1.5 times the interquartile range (IQR) (**D**), and data in (**E**) are presented as mean ± SEM. P values were estimated using two-sided Wilcoxon rank-sum tests with FDR adjustment (**D, E**) or two-way PERMANOVA (**F**).

### Low Biomass and Contamination

In our analyses, we noted that the intrinsic microbiome of pancreatic tissue and cyst fluid was extremely low in microbial biomass. For example, pancreatic tissue as an aggregate had a median of 3,682 (range 399-8,979) copies of bacterial DNA/ng total DNA, which was minimally above the copies of bacterial DNA found in our negative control (PBS; median 2,766, range 768-3,432). In contrast, our positive control (saliva) had 290,184 (range 226,082-1,064,923) copies of bacterial DNA/ng total DNA which is over 75 times the bacterial content of the pancreatic tissue (Supplemental Figure 3). We noted that many of the bacteria phyla and genera detected are not typically bacteria found within the gastrointestinal system and that many have previously been implicated as contaminants in extraction kits. [18] Since the endogenous microbial content of our pancreatic samples is extremely low, contamination is likely to contribute to a non-negligible component of the observed bacteria. [19]

Since our tissue had been procured from two different institutions, we evaluated if differences in institutional procurement practices influenced the microbiome detected and could reveal insights into the effects of the ‘kitome” on the detection of the microbiome within our samples. Although we found no difference in overall abundance by procurement institution, bacterial composition differed in samples procured from MSK versus Duke (Bacteriodetes12.8% vs 6.0%, p=0.011; Proteobacteria 80.1% vs 86.5%, p=0.038; *Nitrobacter* 5.6% vs 0.4%, p=0.00; *Stenotrophomonas* 0.0% vs 0.1%, p=0.015). Additionally, beta diversity did show significant clustering by procurement location within both the pancreatic tissue and cyst fluid (Figure 4). This suggests that differences in procurement and tissue handling were associated with bacterial diversity and could account for slight differences in the bacterial composition we saw across our analysis.

**Figure 4:**
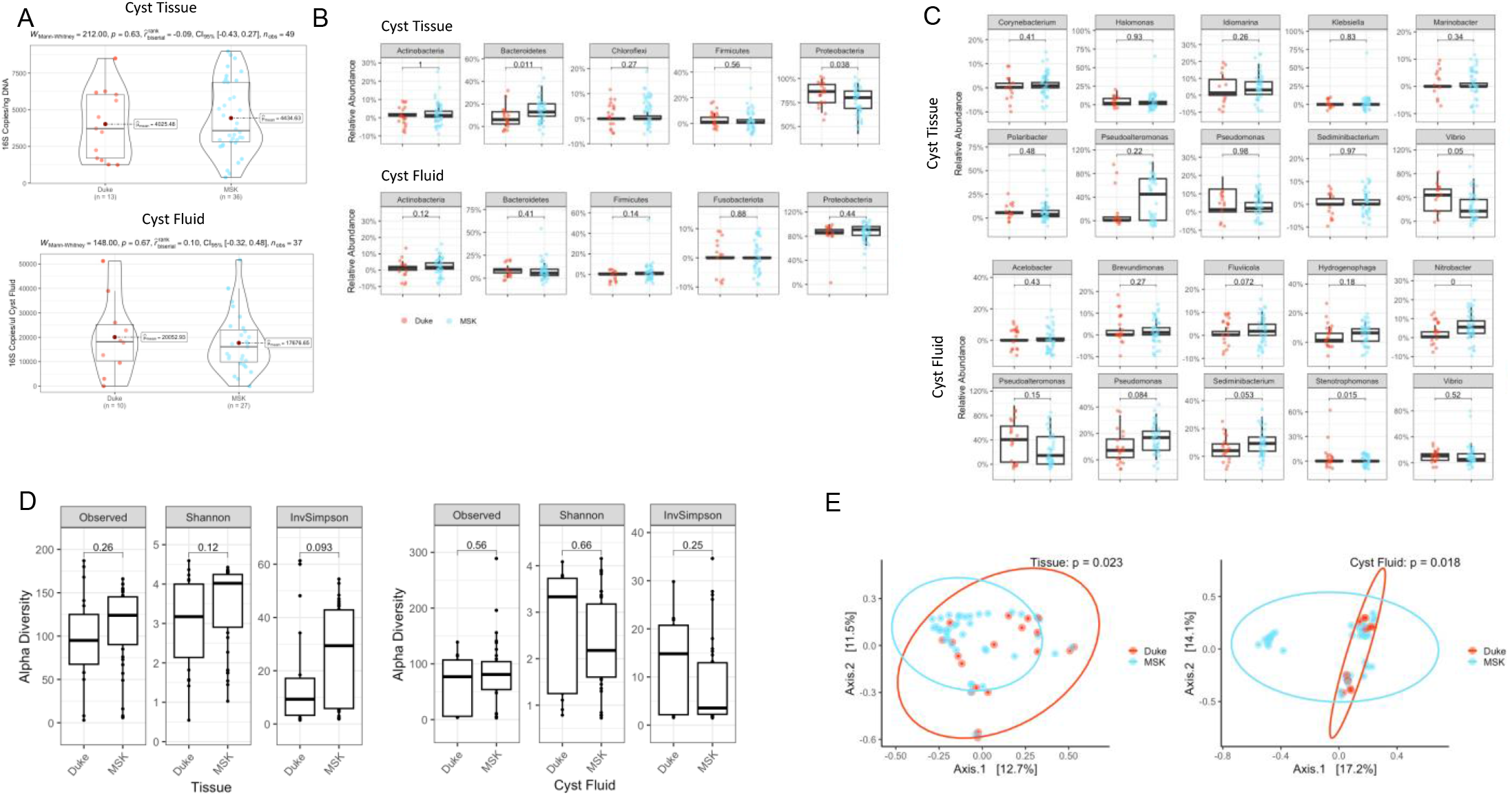
Comparing Pancreatic Microbiome from Samples Procured at Duke vs MSK. A) Absolute read counts of 16S copies/ng of DNA as determined by qPCR in total pancreatic tissue procured from Duke vs MSK and absolute read counts of 16S copies/ul of cyst fluid as determined by qPCR in total pancreatic cyst fluid procured from Duke vs MSK. The mean is marked with a red dot and the median with a horizontal bar. No statistically significant difference was found in the 16S copies/ng DNA between pancreatic tissue or 16S copies/ul of cyst fluid procured from Duke vs MSK as determined by a Mann-Whitney test. B) Box plots of relative abundance at the phyla level between pancreatic tissue and cyst fluid procured at Duke vs MSK. C) Box plots of relative abundance at the genera level between pancreatic tissue and cyst fluid procured at Duke vs MSK. D) Alpha diversity as measured by Observed ASVs, Shannon, and InvSimpson indices between pancreatic tissue and cyst fluid procured at Duke vs MSK. E) PCoA plots of bacteria communities in pancreatic tissue and cyst fluid procured from Duke vs MSK based on the raw Bray-Curtis dissimilarity matrix. Box plot minima and maxima bounds represent the 25^th^ and 75^th^ percentiles, respectively; the center bound represents the median (**B, C, D**). Whiskers extend to 1.5 times the interquartile range (IQR) (**B, C**), and data in (**D**) are presented as mean ± SEM. P values were estimated using two-sided Wilcoxon rank-sum tests (**B, C, D**) or two-way PERMANOVA (**E**).

### CPTAC Analysis

Given that we noted differences in the microbiome composition from samples procured at different institutions, we sought to identify and remove contamination to accurately define the intrinsic pancreatic microbiome. We hypothesized that if a true microbiome exists within the pancreas, the bacteria identified in this study could be confirmed independently using an external cohort of sequenced pancreatic tissue. Our decontamination strategy removed 99.3% of all genera detected in CPTAC sequencing data, leaving 12 genera whose presence in pancreatic tissue could not be explained by contamination (Figure 5). Most of these candidate bacteria are known to colonize the lower GI tract.

**Figure 5:**
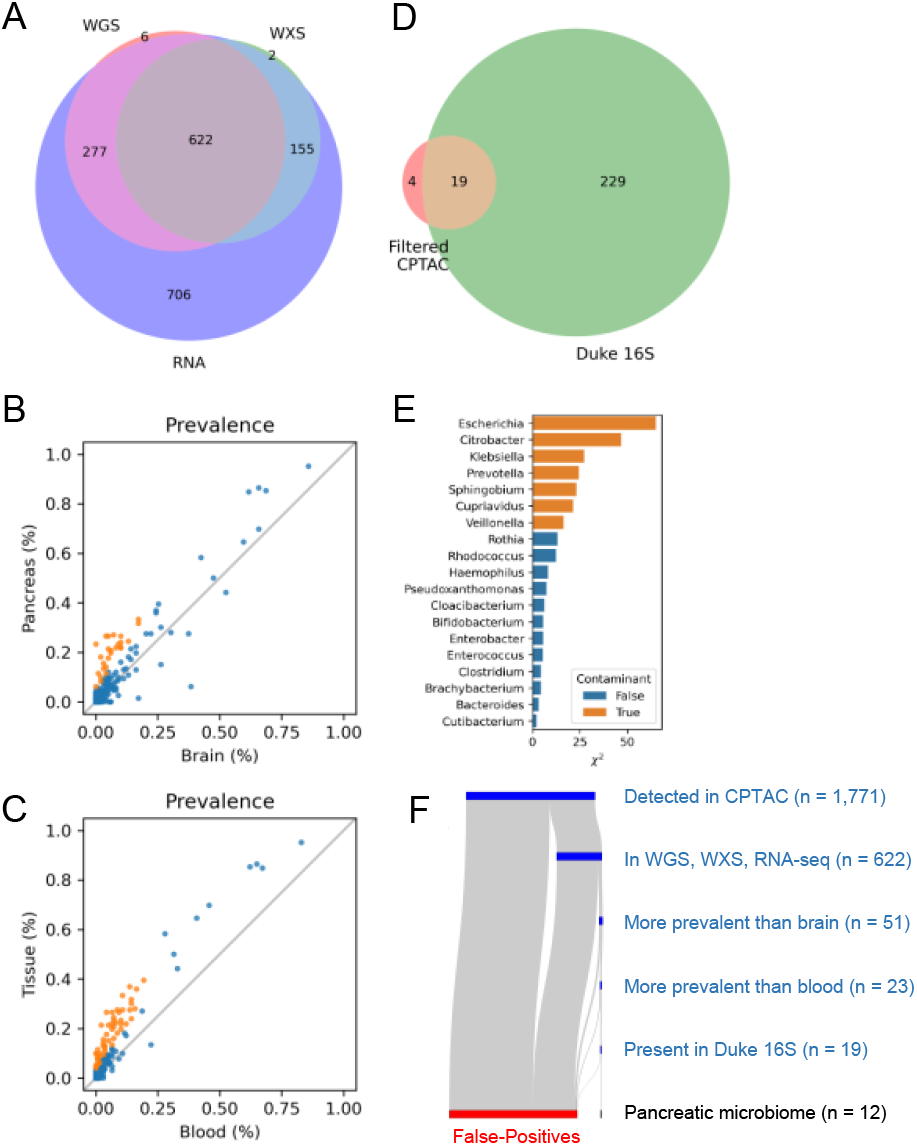
Decontamination Processes using CPTAC External Database. A) A total of 1,771 genera were identified in the 140 CPTAC pancreas samples. There were 622 common genera detectible in all three sequencing assays (WGS, WXS, and RNA-seq) B) Prevalence analysis comparing bacteria genera found in pancreatic tissue to sterile brain tissue. Genera were classified as potentially tissue-resident if they were more prevalent in pancreatic tissue (p < 0.05) and were detected in fewer than 20% of brain samples. C) Prevalence analysis comparing bacteria genera found in pancreatic tissue to matched blood. Genera were classified as potentially tissue-resident if they were more prevalent in pancreatic tissue (p < 0.05) and were detected in fewer than 20% of blood or brain samples. Only 23 genera met these requirements. D) Of the resulting 23 genera that passed frequency and prevalence analysis, only 19 were detected within a second PDAC cohort (Duke). E) A Chi-Square test was used to determine if the observed frequency at which genera were detected in each batch differed from the expectation. Seven of the 19 genera were found to be contaminants, leaving only 12 genera whose presence in pancreatic tissues could not be explained by contamination, false-positive alignment, or batch effect. F) Flow diagram of decontamination pipeline. This decontamination strategy removed 99.3% of all genera detected in CPTAC sequencing data, leaving a total of 12 genera whose presence in pancreatic tissue could not be explained by contamination.

After defining the 12 genera that appeared to be intrinsic to the pancreas after decontamination, we compared them across disease states in our 16S rRNA gene sequencing data. There was no difference in composition, alpha diversity, or beta diversity in the tissue between normal controls, IPMN, and PDAC, likely due to their low overall abundance. Of the 12 intrinsic bacteria genera, only *Psuedoxanthomonas* had greater abundance in the cyst fluid of HGD IPMN than the LGD IPMN (p=0.017). However, there was no difference in *Psuedoxanthomonas* abundance in pancreatic tissue. We found no other differences in composition in the tissue or cyst fluid between LGD and HGD IPMNs and no difference in alpha or beta diversity (Figure 6).

**Figure 6:**
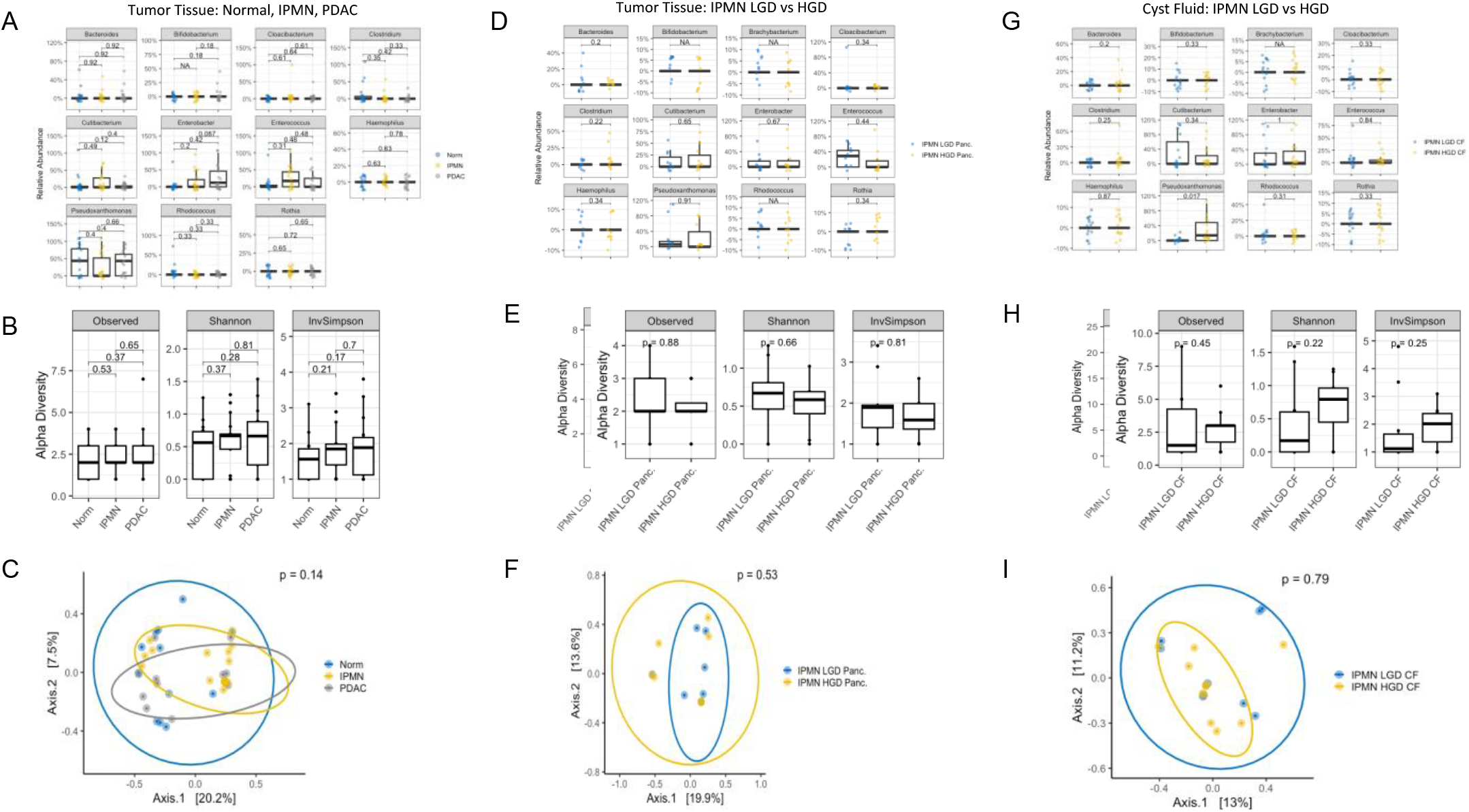
Decontaminated Intrinsic Pancreatic Microbiome Across Diseased States. A) Box plots of relative abundance at the genera level between normal pancreatic tissue, IPMN, and PDAC. B) Alpha diversity as measured by Observed ASVs, Shannon, and InvSimpson indices between normal pancreatic tissue, IPMN, and PDAC. C) PCoA plot of bacteria communities in normal pancreatic tissue, IPMN, and PDAC based on the raw Bray-Curtis dissimilarity matrix. D) Box plots of relative abundance at the genera level between IPMN LGD and HGD pancreatic tissue. E) Alpha diversity as measured by Observed ASVs, Shannon, and InvSimpson indices between IPMN LGD and HGD pancreatic tissue. F) PCoA plot of bacteria communities in IPMN LGD versus IPMN HGD pancreatic tissue based on the raw Bray-Curtis dissimilarity matrix. G) Box plots of relative abundance at the genera level between IPMN LGD and HGD cyst fluid. E) Alpha diversity as measured by Observed ASVs, Shannon, and InvSimpson indices between IPMN LGD and HGD cyst fluid. F) PCoA plot of bacteria communities in IPMN LGD versus IPMN HGD cyst fluid based on the raw Bray-Curtis dissimilarity matrix. Box plot minima and maxima bounds represent the 25th and 75th percentiles, respectively; the center bound represents the median (**A, B, D, E, G, H**). Whiskers extend to 1.5 times the interquartile range (IQR) (**A, D, G**), and data in (**B, E, H**) are presented as mean ± SEM. P values were estimated using two-sided Wilcoxon rank-sum tests with FDR correction (**A, B, D, E, G, H**) or two-way PERMANOVA (**C, F, I**).

Further, the CPTAC database contains 140 pancreatic tumor samples and matched adjacent normal pancreatic tissue. We compared relative abundances of the 12 tissue-resident bacteria between the CPTAC tumor and CPTAC adjacent normal tissue. *Rhodococcus* was the only bacterial genera with differential abundance and had greater abundance in normal tissue than tumor tissue (p=0.003) (Supplemental Figure 3).

## Discussion

Our findings demonstrated that both pancreatic tissue and cyst fluid have a low presence of microbial DNA that does not appear to differ between normal pancreas, IPMN, or PDAC. Across tissue types, there were no statistically significant differences in bacteria composition or diversity. Similarly, the cyst fluid analysis showed extremely limited differences in the microbial composition and did not show a difference in alpha or beta diversity.

The extremely low biomass of these samples raised concerns that a significant portion of the detected microbiota resulted from contamination. Next generation sequencing (NGS) amplifies intrinsic and contamination-associated bacteria equally, meaning that in lower biomass samples, it is increasingly difficult to distinguish the intrinsic microbiome from contamination [18–20]. Prior work has established a lower limit (10^6^ bacteria/ml) where bacterial detection by NGS is no longer considered accurate. [21] In our study, we determined that the bacterial density within pancreatic cyst fluid and tissue falls just at the lower threshold of 10^6^ bacteria/ml, and thus microbiome composition should be interpreted with caution.

Given the low level of microbial DNA detected, we turned to an external dataset and prevalence/validation studies to explore the likelihood of contamination. We used publicly available CPTAC sequencing data with matched PDAC tissue, adjacent normal tissue, and blood samples. Sterile brain tissue from CPTAC was also used for the comparative analysis. If a bacterial genus was more prevalent in pancreatic tissue than blood or brain tissue and was detected by three separate methods (WGS, WXS, and RNA-seq), we concluded it was tissue-resident. We then compared decontaminated bacterial populations from CPTAC to the bacteria found within our samples. There were 12 overlapping bacterial genera that we believe may provide evidence of true tissue resident organisms. However, when we analyzed if these 12 intrinsic bacteria differed across disease states within our original cohort, we did not see a difference in composition or diversity of these bacteria across pancreatic tissue types (normal, IPMN, PDAC). Thus, we conclude that there may be a low biomass, intrinsic microbial presence in the pancreas, but there was no evidence that these organisms differed across disease states.

Prior studies have reported discordant results regarding whether the microbiome is altered in PDAC compared to the normal pancreas. [3, 6–8, 22–24] The majority of these studies rely on NGS, which equally amplifies bacterially derived nucleic acids i and contaminants. As advances in NGS approaches have allowed greater sensitivity, differentiating between contamination and a true bacterial signature has become increasingly difficult. [25] We recently evaluated the publicly available sequencing data from a paper reporting an increased presence of Malassezia in pancreatic tissue from patients with pancreatic cancer relative to normal pancreata. [3] This observation was utilized as the basis for mouse modeling experiments which suggested that the presence of Malassezia in the pancreas may cause pancreatic cancer progression in humans.

Utilizing a panel of computational approaches, our analysis of these data identified only 17 total reads of Malassezia within the 13 pancreatic cancer tissues, all of which belonged to a single sample. [9]

Others have also reported an abnormal intrinsic pancreatic microbiome in the setting of pancreatic cancer. Pushalkar et al. performed 16S rRNA sequencing on PDAC tumors from 12 patients and found that the bacterial composition in human PDAC was distinct from that of the normal human pancreas. [6] However, three of the genera that Pushalkar defined as most abundant and prevalent in human PDAC tissue have been previously identified as common contaminants found in DNA extraction blank controls. [20, 25–27] In contrast, Del Castillov et al. performed 16S rRNA sequencing on both tissue and swabs from multiple gastrointestinal locations (pancreas, bile duct, duodenum, stomach) from 116 patients with PDAC or other conditions requiring foregut surgery. They noted that the bacterial signatures detected within the pancreas were subject-specific rather than disease-specific, consistent with our conclusion that we currently do not have evidence of differing microbial presence by disease state. [22]

Fewer studies have specifically examined IPMNs and whether tissue or cyst fluid microbiome drives dysplastic progression within IPMNs. Gaiser et al. performed 16S PCR and 16S rRNA sequencing on 105 patients with pancreatic cysts and found that a higher 16S bacterial DNA copy number was associated with HGD. [28] In a subsequent study, they cultured cyst fluid obtained during surgical resection and found that 24% of cases exhibited bacteria growth and that a greater portion of patients with HGD IPMN vs LGD IPMN were in the culture-positive cohort.[29] Of note, in both of their studies, 16S read counts and culture positivity were associated with invasive endoscopy, which could be a source of bacterial introduction and contamination. [28, 29] We performed a subset analysis stratifying patients by invasive endoscopy and found no bacterial abundance or compositional differences in patients with IPMN. Importantly, prior IPMN microbiome studies have focused solely on cyst fluid, whereas we analyzed both cyst fluid and tissue. These analyses consistently demonstrate extremely limited differences in the microbial composition as IPMN dysplasia progresses. A general limitation of pancreatic microbiome research, and in our study, is defining control tissue that represents a true normal pancreas. In this study, our control tissue was histologically normal pancreatic tissue adjacent to small, low-grade PNET, but we acknowledge that PNET tumors and surrounding tissue may contain microbiome alterations. Additionally, patients diagnosed with pancreatic cancer vs pancreatic cysts are often in different overall physical conditions, may consume different diets, and may be exposed to different interventions (e.g., chemotherapy, biliary stenting). All have the potential to alter their gut microbiome and, thus, potentially, their intrinsic pancreatic microbiome. Nonetheless, this study’s strength lies in its high number of IPMN and PDAC tumor samples, analysis of both pancreatic tissue and cyst fluid, and careful attention to decontamination in low biomass samples.

In summary, our data suggest that the human pancreas may contain a sparse tissue and cyst fluid microbial population whose composition does not differ across disease states or as malignant progression occurs from LGD IPMN to PDAC. Advances in next-generation sequencing technology have allowed identification of bacterial species in organ spaces with low bacterial burden. However, the increased sensitivity of next-generation sequencing increases the challenge of defining a true microbiome signature versus contamination and calls for increased discrimination in interpreting low-biomass results.

## Acknowledgments

We acknowledge the Duke University BioRepository & Precision Pathology Center (Duke BRPC; supported by P30CA014236) and the National Cancer Institute’s Cooperative Human Tissue Network (CHTN; supported at Duke University by UM1CA239755) for their regulatory and technical assistance in the procurement of the patient samples utilized in this study. Additionally, we would like to acknowledge the UNC microbiome core for their assistance in analyzing our data and CPTAC for providing access to their database and metadata. Data used in this publication were generated by the Clinical Proteomic Tumor Analysis Consortium (NCI/NIH).

