## Supplemental Figures for "Comprehensive Assessment of the Intrinsic Pancreatic Microbiome"

### Supplemental Figure 1- Comparing the Microbiome of Instrumented vs Non-instrumented Pancreatic Cysts

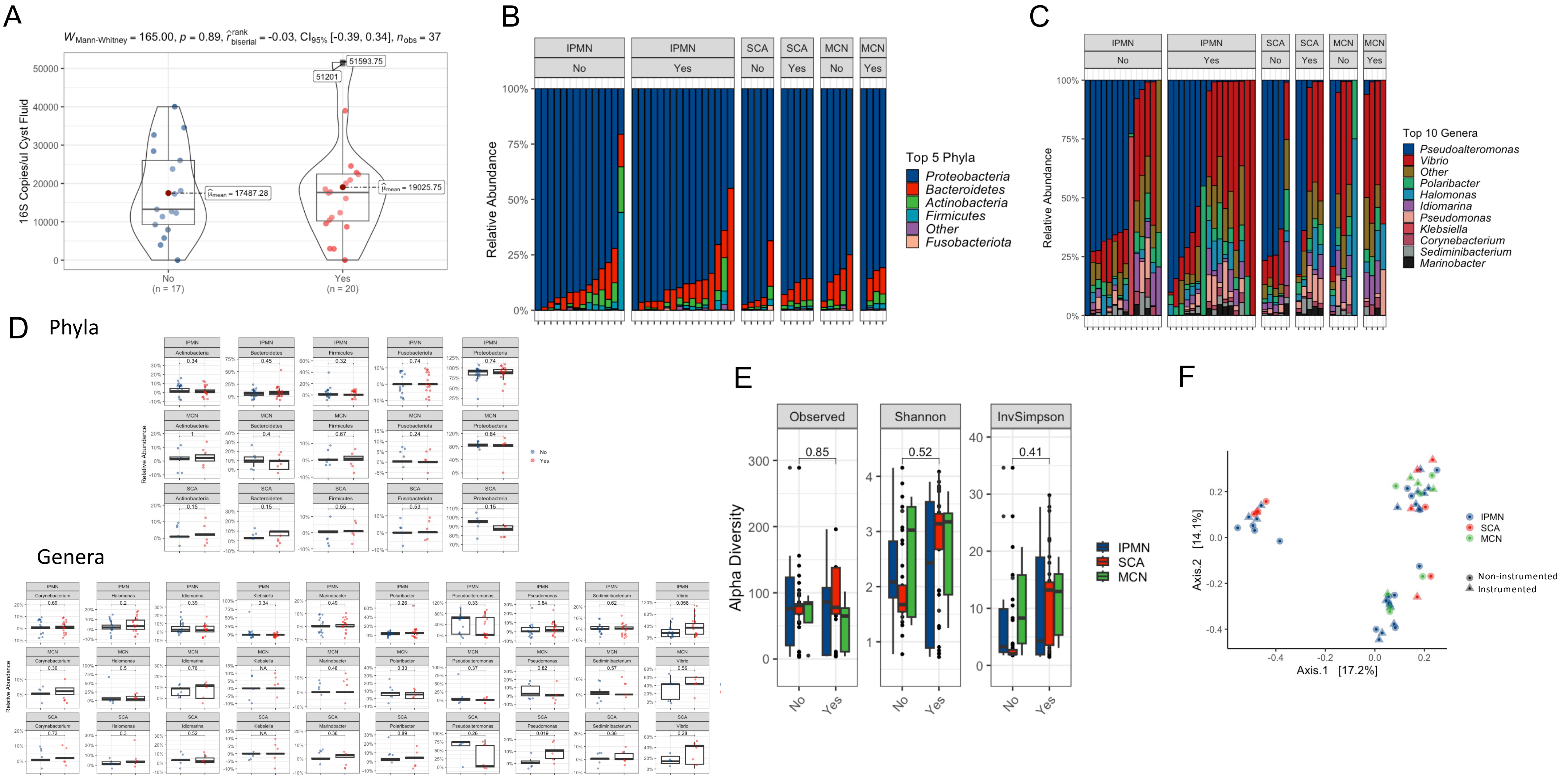

### Supplemental Figure 2: Patients Receiving Antibiotics within Three Months prior to Surgery

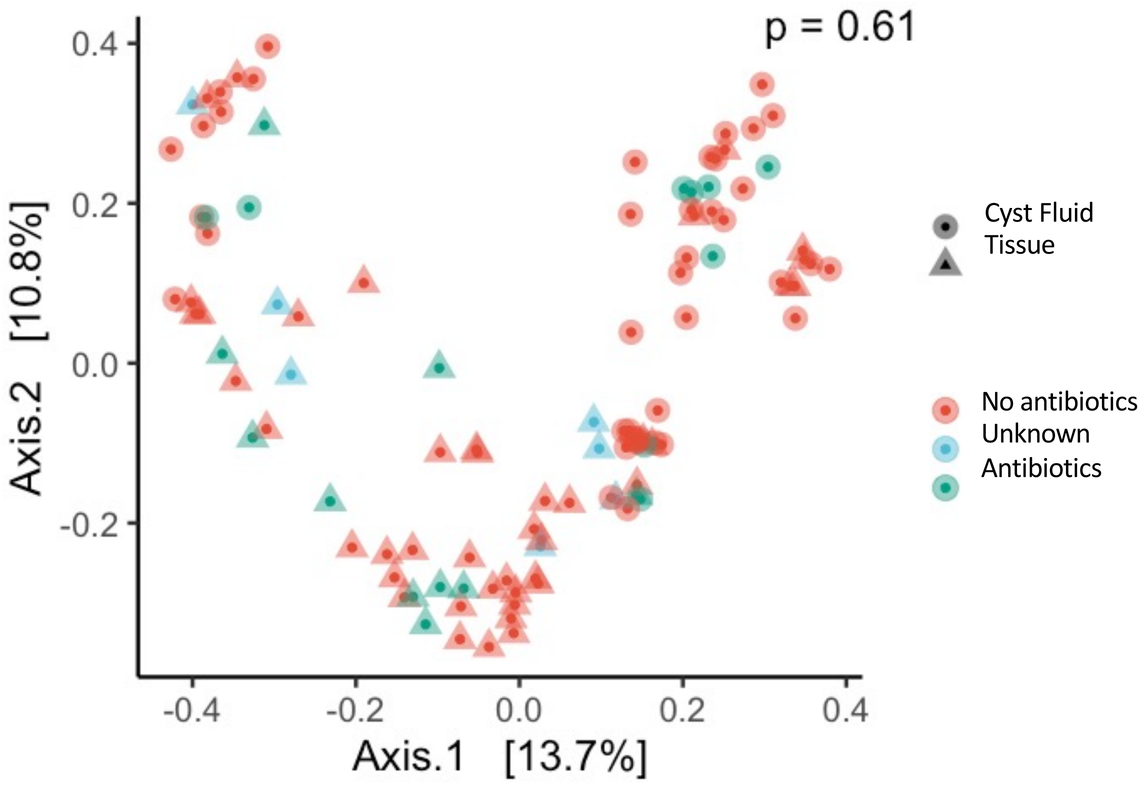

Supplemental Figure 3: Read Counts Across Tissue/Cyst Fluid and Controls

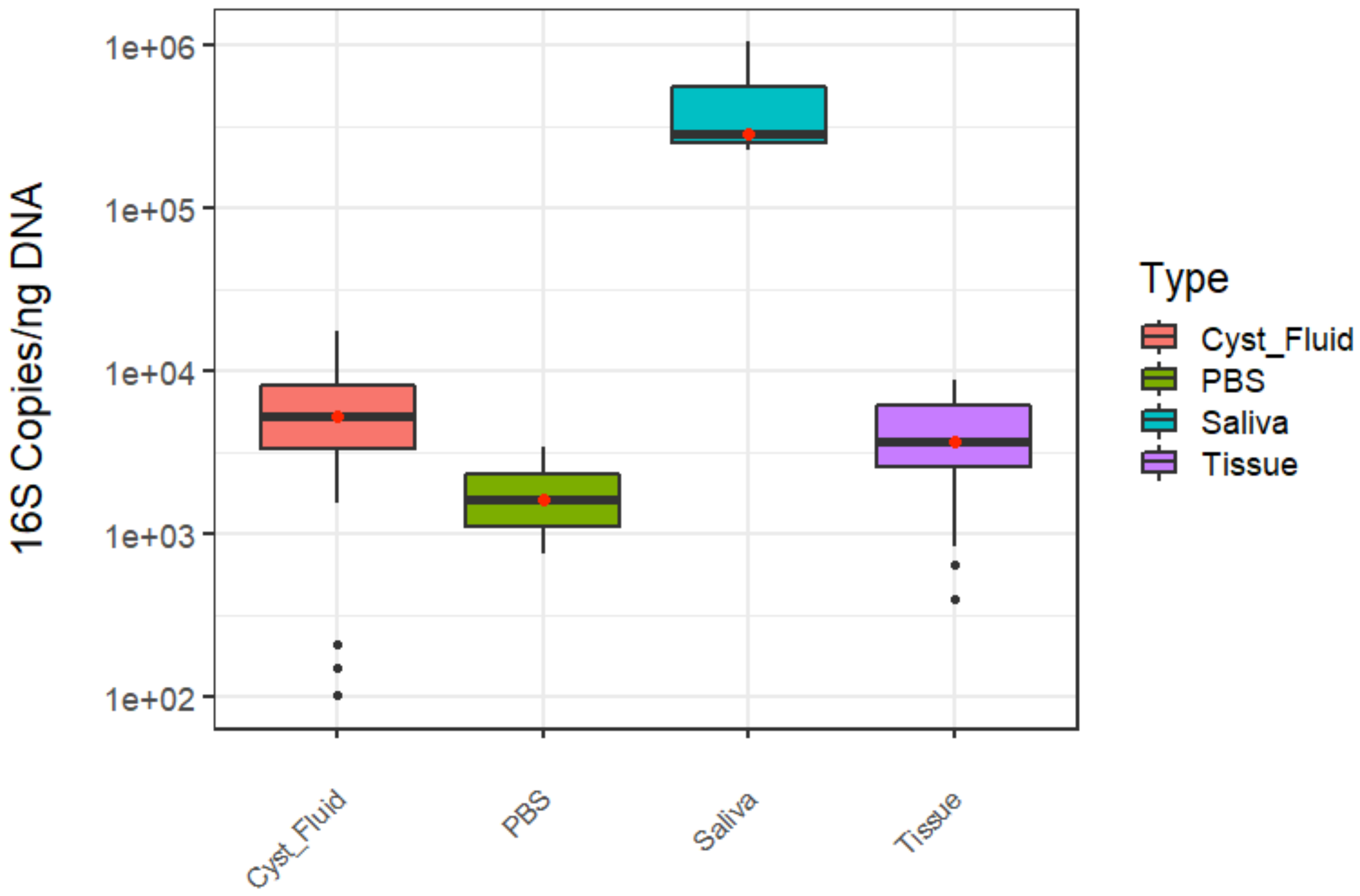

### Supplemental Figure 4: CPTAC Data – Comparing Tumor Tissue to Normal Adjacent Tissue

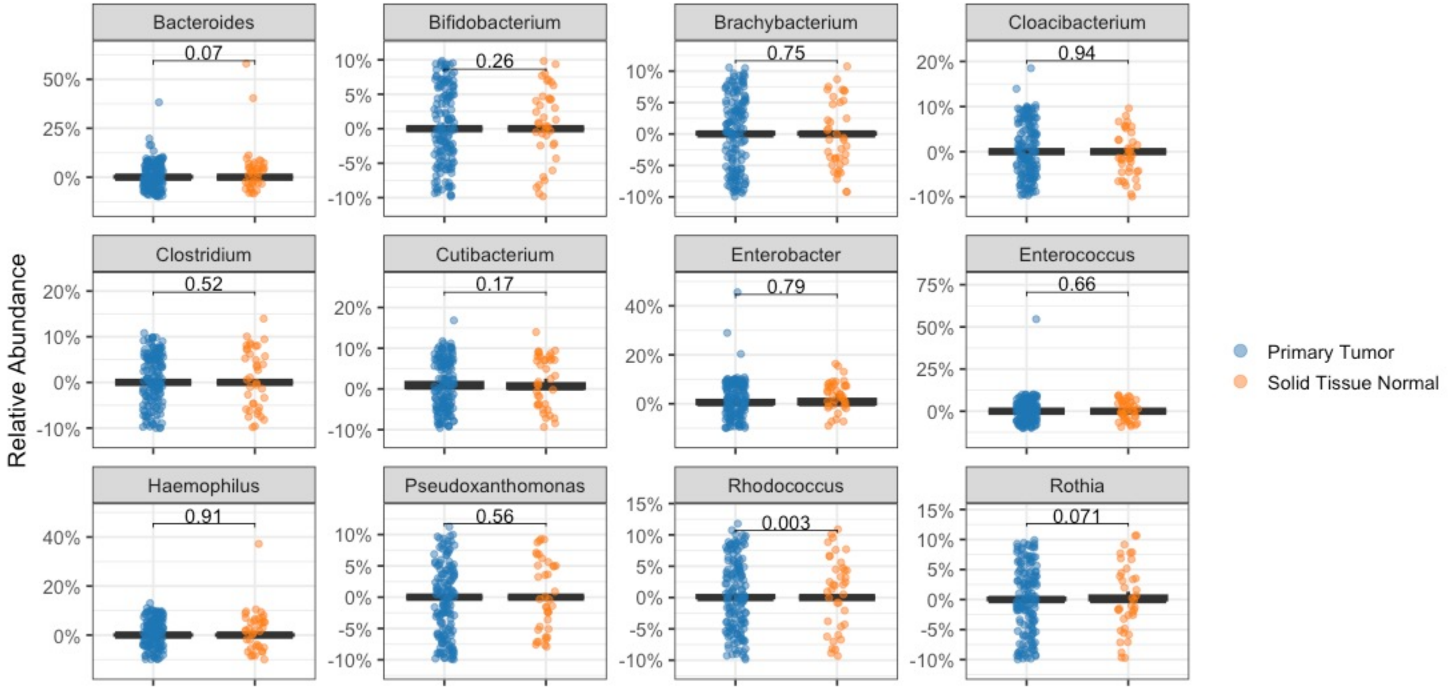


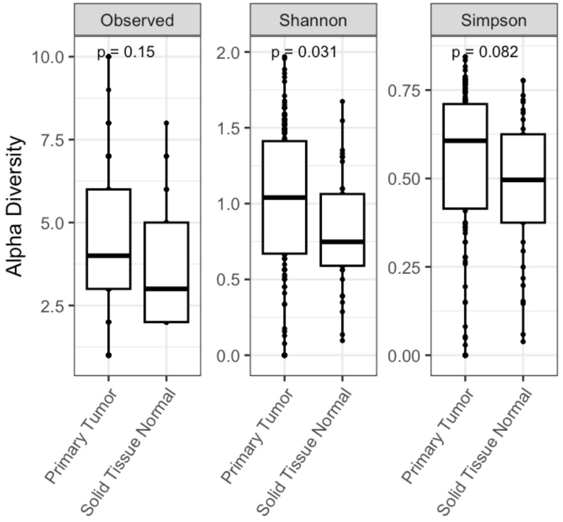


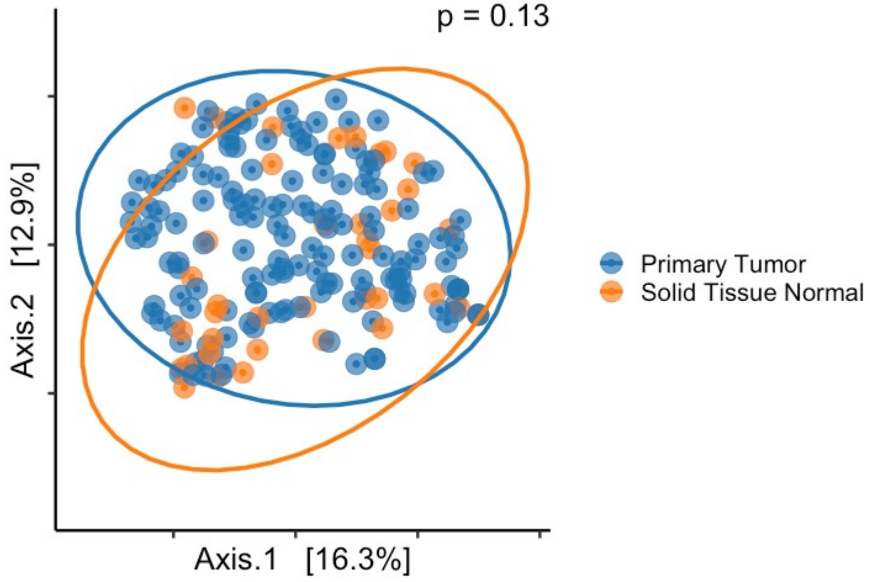
